# Systematic screening of tryptophan metabolism identifies site- and microbial-specific signatures of tryptophan utilization in experimental colitis

**DOI:** 10.64898/2025.12.16.693850

**Authors:** Lina Welz, Abrar I Alsaadi, Danielle MM Harris, Meiping Yu, Taous Mekdoud, Johanna Bornhäuser, Eva Springer, Clara Gilloteau, Anant A. Pothakamury, Jaclyn D. Smith, Brenita C. Jenkins, Felix Sommer, Silvio Waschina, Philip Rosenstiel, Stefan Schreiber, Melanie R McReynolds, Konrad Aden

**Author notes:** Contributed equally (Lina Welz, Abrar I Alsaadi). Shared senior authorship (Melanie R McReynolds, Konrad Aden). Correspondence to Konrad Aden, MD Department of Internal Medicine I, University Hospital Schleswig Holstein Arnold-Heller-Str. 3, 24105 Kiel, Germany.

## Abstract

**Background:** Altered tryptophan (Trp) metabolism and disrupted nicotinamide adenine dinucleotide (NAD⁺) synthesis are hallmarks of IBD, yet how intestinal microbiota contribute to these metabolic shifts during intestinal inflammation remains poorly understood.

**Methods:** We used targeted metabolomics to systematically profile Trp- and NAD⁺-related metabolites across multiple biological compartments – including tissues, luminal contents, stool, and serum – in mice treated with dextran sulfate sodium (DSS) alone or in combination with a broad-spectrum antibiotic (ABX) cocktail.

**Results:** Microbial depletion significantly attenuated colitis and increased host Trp bioavailability, implicating the gut microbiota as a competitive Trp consumer. In DSS colitis, Trp degradation along the kynurenine pathway (KP) was exaggerated but blocked at the key KP enzyme quinolinate phosphoribosyltransferase (QPRT), resulting in mucosal NAD(H) depletion. ABX co-treatment normalized metabolite conversion along the KP and restored mucosal NAD(H) levels, revealing a dual role of the gut microbiota during colitis: while they compete with the host for Trp utilization, they simultaneously shape host KP regulation and NAD⁺ *de novo* synthesis, supporting host energy homeostasis.

**Conclusion:** Our findings demonstrate that mucosal NAD⁺ *de novo* synthesis is a microbially regulated metabolic process that alleviates intestinal inflammation and may represent a novel therapeutic target in IBD through modulation of the gut microbiota or their metabolites.

## Introduction

The gut microbiota play a central role in the pathogenesis of inflammatory bowel disease (IBD), a chronic-relapsing inflammatory disorder affecting the intestine comprising two main sub entities, Crohn’s disease (CD) and ulcerative colitis (UC). Alterations in microbial composition and function including reduced diversity, increase of pathobionts and loss of protective bacteria, often termed dysbiosis, are a hallmark of IBD [1–3]. These community shifts reflect uncoupled dynamics between the intestinal microbiota and the host, which are not only observed at the functional level, but also exhibit temporal variability, becoming more predominant upon active disease and reflecting compromised microbial resilience [4,5]. The resulting loss of ecological stability contributes to chronic intestinal inflammation by impairing epithelial barrier integrity, increasing luminal translocation, fuelling aberrant immune responses, and perturbing host metabolism [6–10]. Previous studies using germ-free (GF) mouse models, murine models of microbial depletion by antibiotics and microbiota-transfer experiments have established that the composition of the gut microbiota and its interplay with the host influence the development of colitis [11–16].

Tryptophan (Trp) metabolism has emerged as a central intersection between host-microbial crosstalk and the pathogenic mechanisms underlying IBD [17,18]. The majority of intestinally absorbed Trp (>90%) is degraded along the kynurenine pathway (KP), which in the gut is exerted by indoleamine 2,3-dioxygenase 1 (IDO1) and feeds into nicotinamide adenine dinucleotide (NAD^+^) *de novo* synthesis, whereas the indole pathway, exerted by the gut microbiota, and the serotonin pathway, executed by enterochromaffin cells, to minor extents contribute to Trp turnover [19]. Importantly, the final product of the KP, NAD^+^, is an essential redox co-enzyme which regulates cellular energy supply, immune responses, DNA repair and metabolism [20,21]. Next to *de novo* synthesis from Trp, NAD^+^ is synthesized via the salvage pathway (nicotinamide (NAM), nicotinamide riboside (NR)) or the Preiss-Handler pathway (from nicotinic acid (NA)) [22–24]. Exaggerated turnover of Trp along the KP has been identified as an overarching hallmark of chronic inflammatory diseases, including IBD [25,26]. We have previously shown that despite increased Trp catabolism, NAD⁺ *de novo* synthesis is impaired due to a critical bottleneck at the level of the KP enzyme quinolinate phosphoribosyltransferase (QPRT), causing accumulation of the NAD⁺ precursor quinolinic acid (Quin) and insufficient turnover into NAD⁺ [27]. The resulting mucosal shortage of NAD⁺ has been reported to disrupt mitochondrial function and cellular energy supply, fuelling inflammation, whereas the coinciding elevation of serum Quin was observed in individuals who later developed Crohn’s disease, suggesting its potential as a predictive biomarker [28,29]. Importantly, mucosal NAD^+^ exhaustion has been described as a critical pathophysiological event in inflammatory disorders, particularly in IBD [30,31].

Not only inflammatory processes but also gut microbiota affect how ingested Trp is utilized by metabolizing it into various bioactive catabolites. These include indole derivatives such as indole-3-acetic acid (IAA), indole-3-propionic acid (IPA) and tryptamine, while also modulating host-driven metabolism such as the synthesis of kynurenine (Kyn) or serotonin. Many of these metabolites serve as ligands for the aryl hydrocarbon receptor (AhR), thereby influencing key host functions such as immune regulation, intestinal barrier integrity and the gut-brain axis [32,33].

We hypothesized that gut microbiota affect host Trp and NAD^+^ metabolism during intestinal inflammation. To date, a comprehensive, system-wide analysis of how the gut microbiota influences host Trp metabolism during intestinal inflammation is lacking. The aim of our study was therefore to define the impact of the intestinal microbiota on colitis-induced Trp catabolism and dysregulated NAD⁺ metabolism. To test this, we systematically profiled Trp and its downstream metabolites across multiple organs during experimental colitis, using either antibiotics-treated or GF mice to dissect microbiota-dependent effects.

## Materials and methods

### Animal Use and Care

All animal experiments were either performed at the Pennsylvania State University and approved by the Institutional Animal Care and Use Committee (IACUC - PROTO202202188).) or at the Central Animal Facility (ZTH) of the University Hospital Schleswig Holstein (UKSH, Kiel, Germany) approved by the local animal safety review board of the federal ministry of Schleswig Holstein (IX 554 - 62160/2024 (48-6/24) and internal §4 project number 1410). 8-12-week-old C57BL/6J male mice were purchased from The Jackson Laboratory and received laboratory diet 5010 and water *ad libitum* while they acclimated for at least 5 days in a specific pathogen-free facility. Animals were exposed to a 12-hour-light-dark cycle (7 a.m.-7 p.m.) during experimental use. GF and specific-pathogen free (CONV) mice were housed in the Central Animal Facility (ZTH) of the University Hospital Schleswig Holstein (UKSH, Kiel, Germany) in individually ventilated cages (Green Line, Techniplast) or under sterile conditions in gnotobiotic flexible film isolators. All mice were kept under a 12-h light cycle and fed a regular gamma-irradiated chow diet *ad libitum*.

### Acute DSS colitis and Antibiotic Treatment of Mice

For intestinal microbiota depletion, mice were single-housed and orally treated with drinking water ad libitum containing a cocktail of ampicillin (1g/L, Sigma-Aldrich), vancomycin (0.5g/L, Avantor), neomycin (1g/L, Sigma-Aldrich) and metronidazole (1g/L, Sigma-Aldrich) as previously described for 13 days prior to and during DSS colitis [14,15,34]. To offset the unpalatable taste of the antibiotics, 3% sucrose (VWR) was included in the drinking water of all groups to promote consumption. Consumption of drinking water was monitored daily. For DSS treatment supplied with 2.5% DSS (molecular weight 36-50kDa, MP Biomedical) dissolved in autoclaved drinking water for 5 days, followed by regular drinking water. DSS-containing water was exchanged every other day. Control groups were terminated two days earlier than the DSS-treated groups, with their last recorded observations in body weight and disease activity indices (DAI) carried forward for phenotype classification in comparison to the groups sacrificed later. All animals at both facilities were sacrificed via cervical dislocation without use of any anaesthesia.

### Disease Activity Index Assessment

The severity of colitis was assessed by evaluating the following parameters daily: weight loss (0 points = no weight loss or gain, 1 point = 1-5% weight loss, 2 points = 6-10% weight loss, 3 points = 11-19% weight loss, 4 points = 20-25% weight loss, 5 points = >25% weight loss); stool consistency (0 points = normal and firm, 1 point = very slight change, 2 points = slight change and soft, 3 points = moderate change, 4 points = noticeable change, diarrhoea, 5 points = severe change, runny diarrhoea; bleeding stool (0 points = normal, 1 point = redness of perianal region, 2 points = slightly blood-streaked stool, 3 points = blood-streaked stool, 4 points = marked blood contamination, 5 points = bloody stool); posture (0 points = normal, 1 points = very slight change, 2 points = slightly curved, 3 points = moderate change, 4 points = strongly curved, 5 points = severe change and consistently curved); activity (0 point = normal active, 1 point = very slight change in activity, 2 points = reduced movement and clinging onto the cage, 3 points = moderate change, 4 points = noticeable change and rarely clinging onto the cage, 5 points = severe change, sitting still and no movement); fur (0 points = normal, 1 point = very slight change, 2 points = slightly dirty and scruffy, 3 points = moderate change, 4 points = noticeable change, dirty, and scruffy, 5 points = severe change, very dirty, dull, and scruffy). The DAI was calculated by combining scores of the measured parameters.

### Histopathologic Assessment of Murine Intestinal Tissue

The colon was collected postmortem, flushed with PBS and cut open longitudinally. The distal colon was rolled up as a Swiss roll from the distal to the proximal part, fixed in 2% CMC (sodium carboxymethyl cellulose, diluted in MilliQ; Sigma-Aldrich) and snap frozen in liquid nitrogen. Sections for histopathologic assessment were cut and stained with hematoxylin and eosin (H&E). Histopathologic scoring displays the combined score of inflammatory cell infiltration and tissue damage and was performed in a blinded fashion, as previously described [35].

### Sample Preparation of Serum and Tissues for LC-MS

Serum was thawed on ice before adding -80°C 100% 80:20 methanol:water with a volume of 13.5µl solvent/1µl serum. Samples were vortexed, incubated on dry ice for 10min and centrifuged at 4°C, 16,000g for 25min. Supernatants were used for analysis. Frozen tissues, fecal samples and luminal contents were weighed and grounded with liquid nitrogen in a cryomill (Retsch) at 25Hz for 45s. 40:40:20 acetonitrile:methanol:water with a volume of 40μL solvent / 1mg of tissue was added. Samples were vortexed for 15s, incubated on ice for 10min and centrifuged at 4°C, 16,000g for 30min. Supernatants were transferred to new tubes and again centrifuged at 16,000g for 25min, this step was repeated. Supernatants were transferred to liquid-chromatography mass spectrometry (LC-MS) vials for analysis.

### Targeted LC-MS of Trp and NAD+ Derivatives

The LC-MS method was based on hydrophilic interaction chromatography (HILIC) coupled to the Orbitrap Exploris 240 mass spectrometer (ThermoFisher Scientific) [36] LC-separation was performed on a XBridge BEH Amide column (2.1x150 mm, 3.5µm particle size, Waters). Solvent A: 95%:5% H2O:acetonitrile with 20mM ammonium bicarbonate, solvent B: acetonitrile. Gradients: 0min, 85% B; 2min, 85% B; 3min, 80% B; 5min, 80% B; 6min, 75% B; 7min, 75% B; 8min, 70% B; 9min, 70% B; 10min, 50% B; 12min, 50% B; 13min, 25% B; 16min, 25% B; 18min, 0% B; 23min, 0% B; 24min, 85% B; 30min, 85% B. Other LC parameters: flow rate 150ml/min, column temperature 25°C, injection volume 10μL and autosampler temperature 5°C. The MS was operated in both negative and positive ion mode. Other MS parameters: resolution 140,000 at m/z 200, automatic gain control target 3e6, maximum injection time 30ms and scan range m/z 70-1000. Raw LC-MS data were converted to mzML format using the command line ‘msconvert’ utility [37]. Data were analyzed with the EL-MAVEN software version 12.

### DNA isolation and bacterial quantification

DNA from fecal samples collected from all mouse groups at baseline (i.e., prior to any treatment) as well as on DSS colitis days 1, 3 and 7 was extracted using the Dneasy Power Soil Pro kit (Qiagen) according to the manufacturer’s instructions. Real-time PCR amplification was performed using 16S universal primers (forward: ACTCCTACGGGAGGCAG, reverse: GACTACCAGGGTATCTAATCC) and probe (TGCCAGCAGCCGCGGTAATAC) that target the V3-V4 region of the bacterial 16S rRNA gene. PCR was performed in 10µl volume with 4.5µl of TaqMan Fast Universal PCR Master Mix (AppliedBiosystems), 0.5µl TaqMan assay, 1µl fecal DNA template, and 4µl PCR water. Assays were run on a ViiA 7 real-time PCR system (Applied Biosystems) using the following cycling conditions: 95°C for 20 seconds, followed by 45 cycles of 95°C for 1 second, and 60°C for 20 seconds. All assays were conducted in duplicates with a positive control of bacterial DNA and a negative control of water. The absolute number of 16S gene copies was quantified by comparison with a standard curve generated by serial dilution of *E. coli* 16S rDNA. The total 16S rRNA gene count was normalized by mg of feces.

### RNA isolation, cDNA synthesis and Gene Expression Analysis

Using the RNeasy Kit (Qiagen), mRNA was isolated from different cell types. cDNA was synthesized using Monarch Genomic DNA Purification Kit (New England Biolabs) according to the manufacturer’s protocol. To examine gene expression, cDNA samples were subjected to quantitative reverse transcription polymerase chain reaction using TaqMan assays purchased from Integrated DNA Technologies (for TaqMan IDs and RefSeq numbers, see Supplementary Table 1). Reactions were carried out on the Applied Biosystems 7900HT Fast Real-Time PCR System (Applied Biosystems) and relative transcription levels were determined utilizing β2M or β-actin as housekeeping genes.

**Supplementary Table 1:**
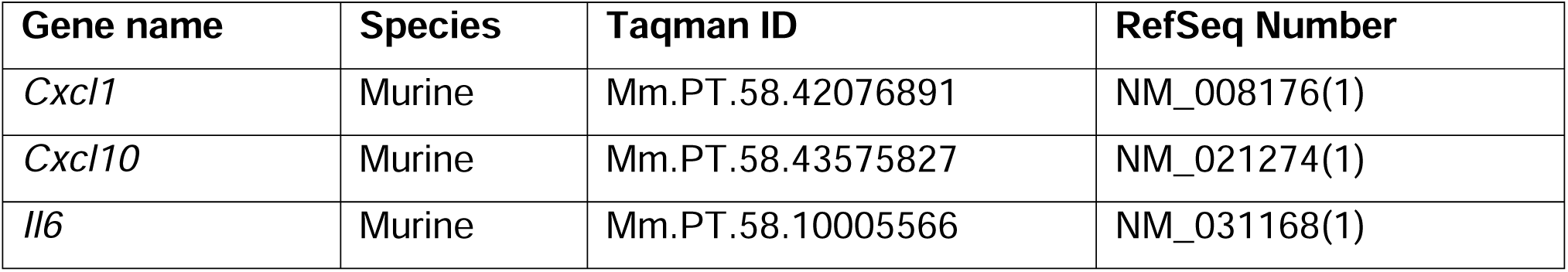
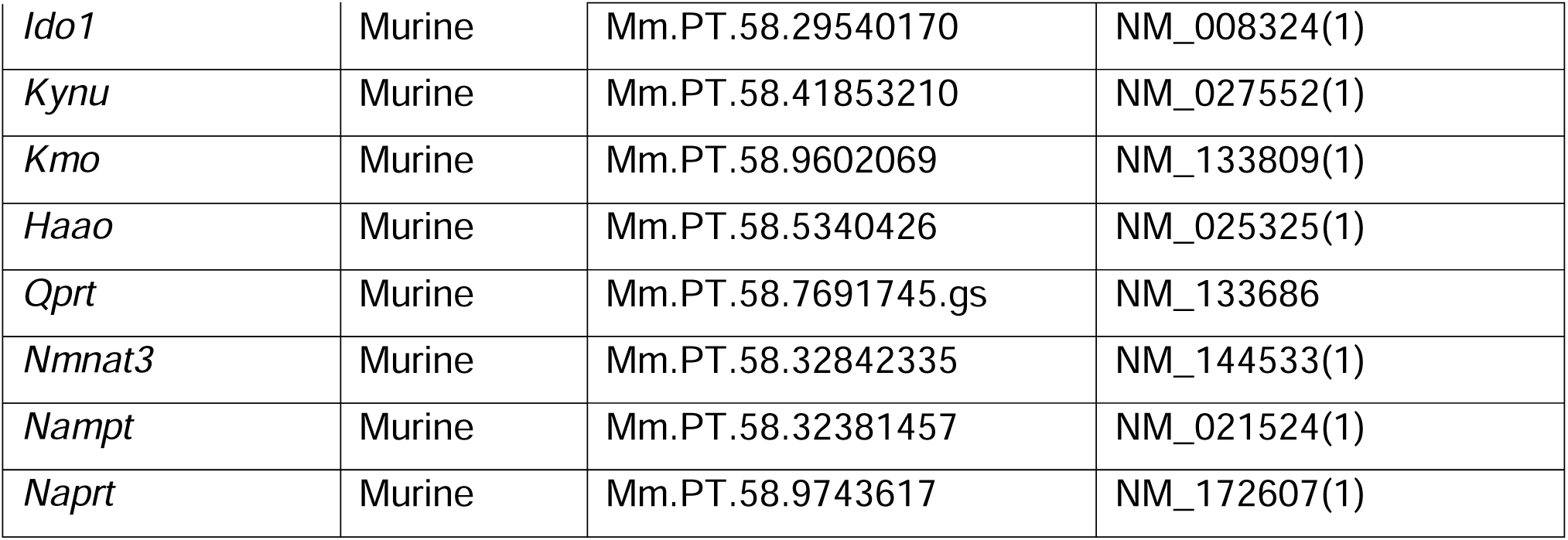
Primers used for gene expression analysis.

### Statistical Analysis

Statistical tests were selected considering data distribution and variance characteristics. Two-way ANOVA followed by Šídák’s or Tukey’s post-hoc test for multiple comparisons were used. A threshold <0.05 was considered statistically significant. Statistical tests were performed with GraphPad PRISM software 10.

## Results

### Microbial depletion protects against DSS colitis

To evaluate how the intestinal influences host Trp metabolism, we depleted gut microbes in young male C57BL/6J mice using a two-week antibiotic cocktail (ampicillin, neomycin, metronidazole, and vancomycin). Following antibiotic pretreatment (ABX), colitis was chemically induced with dextran sodium sulfate (DSS), and antibiotic administration was maintained throughout the experiment (overview in Fig. 1A). To gain insight into systemic metabolic processes, we employed a systematic sampling strategy and, after sacrifice, collected (i) tissues (liver, spleen, kidney, duodenum, jejunum, ileum, cecum, proximal and distal colon), (ii) luminal contents from corresponding gastrointestinal segments, (iii) stool pellets, and (iv) serum (Fig. 1A). The efficacy of microbial depletion following ABX treatment was confirmed by assessing 16S rRNA gene copies in feces of the respective mice, which revealed a 1,000-10,000-fold reduction of the microbial load in ABX-treated animals (Fig. S1A). Microbial depletion was further functionally confirmed by reduced luminal abundances of metabolites strictly depending on microbial synthesis, such as secondary bile acids (lithocholic acid, deoxycholic acid) and the short chain fatty acid (SCFA) butyrate (Fig. S1B-D) [38,39]

**Figure 1:**
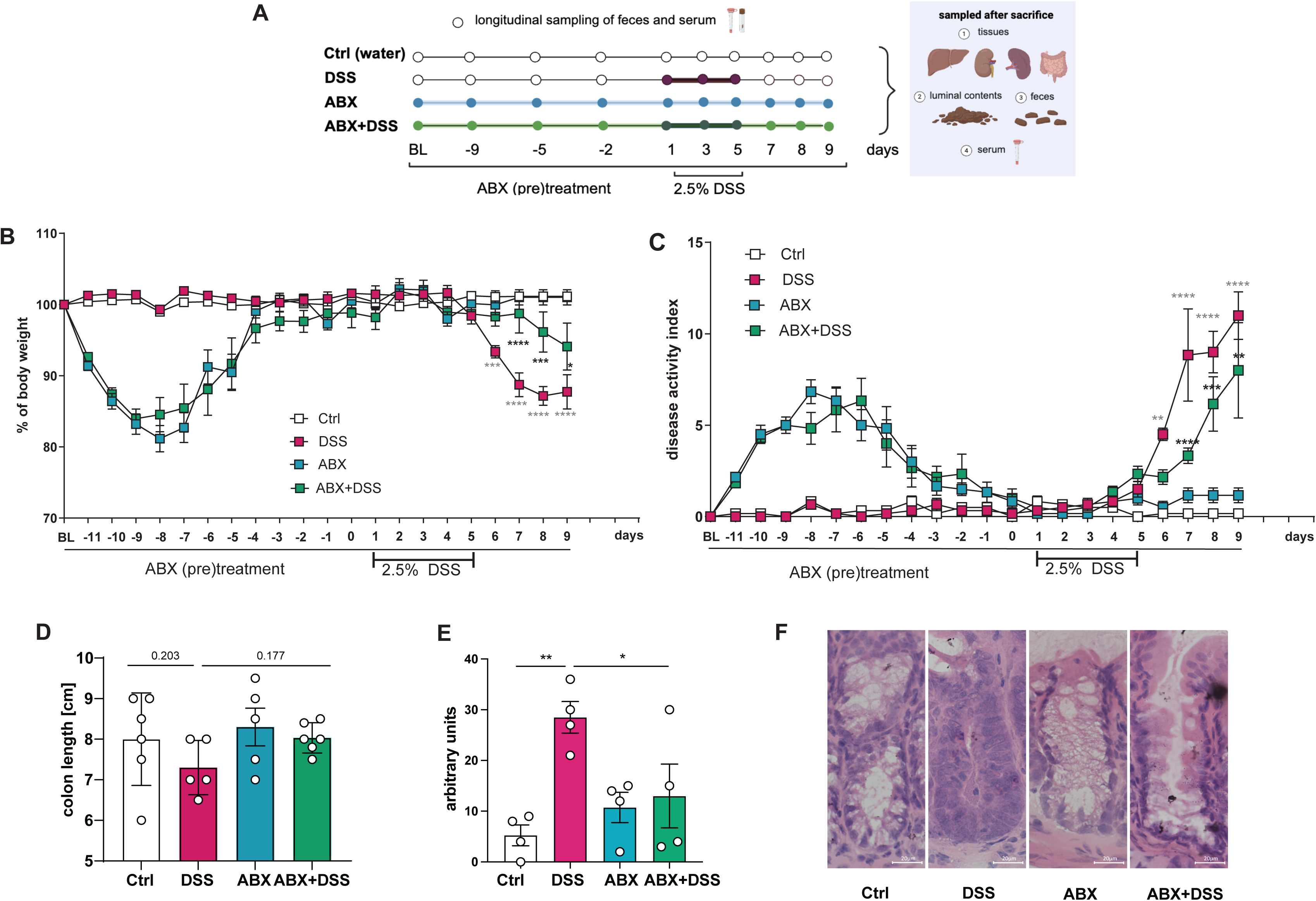
Microbial depletion attenuates severity of DSS-induced colitis. (A) Overview of experimental set-up. 11 weeks old C57BL/6J male mice were pre-treated with an antibiotics (ABX) cocktail for 13 days (blue and green groups), before DSS colitis was induced for 5 days with 2.5% DSS (red and green groups). Sampling of feces and serum is indicated by a round dot by a dot at the respective time points. Post-mortem, luminal contents and tissues were collected. Colitis severity was assessed by (B) body weight loss compared to baseline weight, (C) a combined disease activity index (DAI, comprising relative body weight loss, stool consistency, rectal bleeding, behaviour, activity, fur consistency), (D) colon length, (E) histopathologic assessment with (F) representative images. For (B) and (C), grey asterisks compare control and DSS-treated mice, whereas black asterisks show the difference between DSS and ABX-DSS mice. Data are presented as mean ± SEM. Statistical analysis was performed with two-way-ANOVA. ∗ p<0.05; ∗∗ p<0.01; ∗∗∗ p<0.001, ∗∗∗∗ p<0.0001.

We found that mice receiving ABX were significantly protected from DSS colitis (referred to as “ABX-DSS-mice” in the following), as mirrored by a decreased weight loss and disease activity index score as well as a normalized colon length (Fig. 1B-D). Furthermore, we discovered the colonic histopathological degree of inflammation as well as the expression of pro-inflammatory cytokines of ABX-DSS mice to be strongly reduced in comparison to DSS mice (Fig. 1E,F, Fig. S1E-G).

### Antibiotic treatment influences microbial metabolism during intestinal inflammation

Having shown that microbial depletion alleviates intestinal inflammation, we next sought to determine how the intestinal microbiota shape host Trp metabolism during colitis. We first examined how DSS-induced colitis affects microbial Trp metabolism by quantifying downstream metabolites from major Trp degradation pathways – the kynurenine, serotonin, indole, and pyruvate routes (overview in Fig. S2) – in fecal samples collected on the final day of DSS treatment. We observed increased levels of Trp in ABX-DSS-treated mice, whereas activation of the kynurenine pathway (KP; Kyn:Trp ratio, xanthurenic acid (XanA) and quinaldic acid (QA) was decreased (Fig. 2A-D). In addition, we found that in microbiota-proficient mice, Trp was converted along the serotonin and the indole pyruvate pathway (increased levels of 5-hydroxy indoleacetic acid (5HIAA), IPA and indolelactic acid (ILA)), whereas the respective metabolites were strongly reduced following ABX treatment (Fig. 2E-H). Decreased levels of tryptamine, IAA and indoxyl sulfate were measured both upon DSS, ABX and combined treatment (Fig. S3A-C). In addition, fecal levels of NAD(H) and its precursors – nicotinamide (NAM), nicotinamide riboside (NR), and nicotinic acid (NA) – were reduced in DSS-treated mice, with this decrease further accentuated by microbial depletion (Fig. 2I-M).

**Figure 2:**
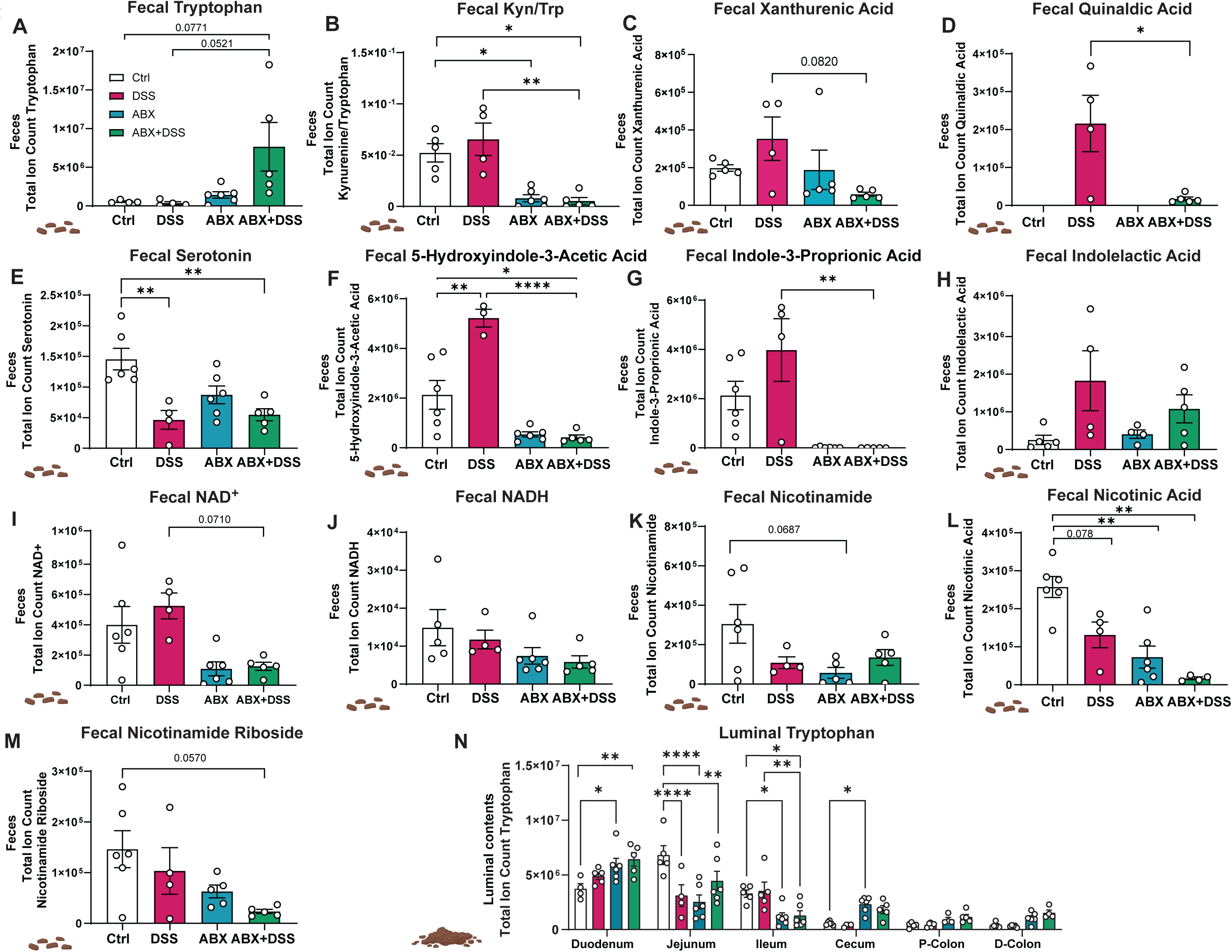
DSS colitis and antibiotics (pre)treatment affect fecal metabolites. Total ion counts of the indicated metabolites are shown in feces (A-M) and for Trp in luminal contents (N). Data are presented as mean ± SEM. Statistical analysis was performed with two-way-ANOVA. ∗ p<0.05; ∗∗ p<0.01; ∗∗∗∗ p<0.0001.

Overall, elevated levels of fecal Trp and depleted conversion into Trp derivatives were observed in ABX-DSS mice, suggesting the intestinal microbiota to act as a substantial consumer of Trp during intestinal inflammation. To determine which regions of the GI tract contributed to the elevated Trp levels, we quantified Trp in luminal contents and found increased abundances in the duodenum and colon (Fig. 2N). We concluded that elevated Trp levels to either result from reduced microbial consumption, decreased mucosal host degradation or a combination of both.

### Increased host Trp bioavailability and reduced shuttling into the KP upon depletion of the intestinal microbiota

Based on the hypothesis that microbial Trp degradation influences microbial and host Trp metabolism, we next measured Trp levels in serum and tissues. Under physiological conditions, Trp was absorbed in the small intestine (SI), where its levels were highest in the jejunum (Fig. 3A). ABX treatment increased Trp availability in the duodenum, cecum and proximal/distal colon. In contrast, ABX treatment reduced mucosal Trp levels in the jejunum and ileum. While DSS treatment caused a non-significant decrease in serum Trp, ABX treatment led to an overall increase in Trp bioavailability in the host (Fig. 3B). In line with normalized Trp levels in ABX-DSS-treated mice, we observed similarly normalized Kyn and Kyn:Trp ratios in serum, colonic tissues and luminal contents from ABX-DSS mice, indicating that excessive turnover of Trp into Kyn in the intestinal mucosa during DSS colitis was ameliorated upon ABX co-treatment (Fig. 3C-F, Fig. S4A,B). Along the same line, *Ido1* expression in the distal colon of ABX-DSS mice was normalized (Fig. S5A). However, appreciable levels of Kyn remained detectable, particularly in the serum of ABX-DSS mice, indicating that Trp-to-Kyn conversion persisted despite the absence of excessive kynurenine pathway activation (Fig. 3D).

**Figure 3:**
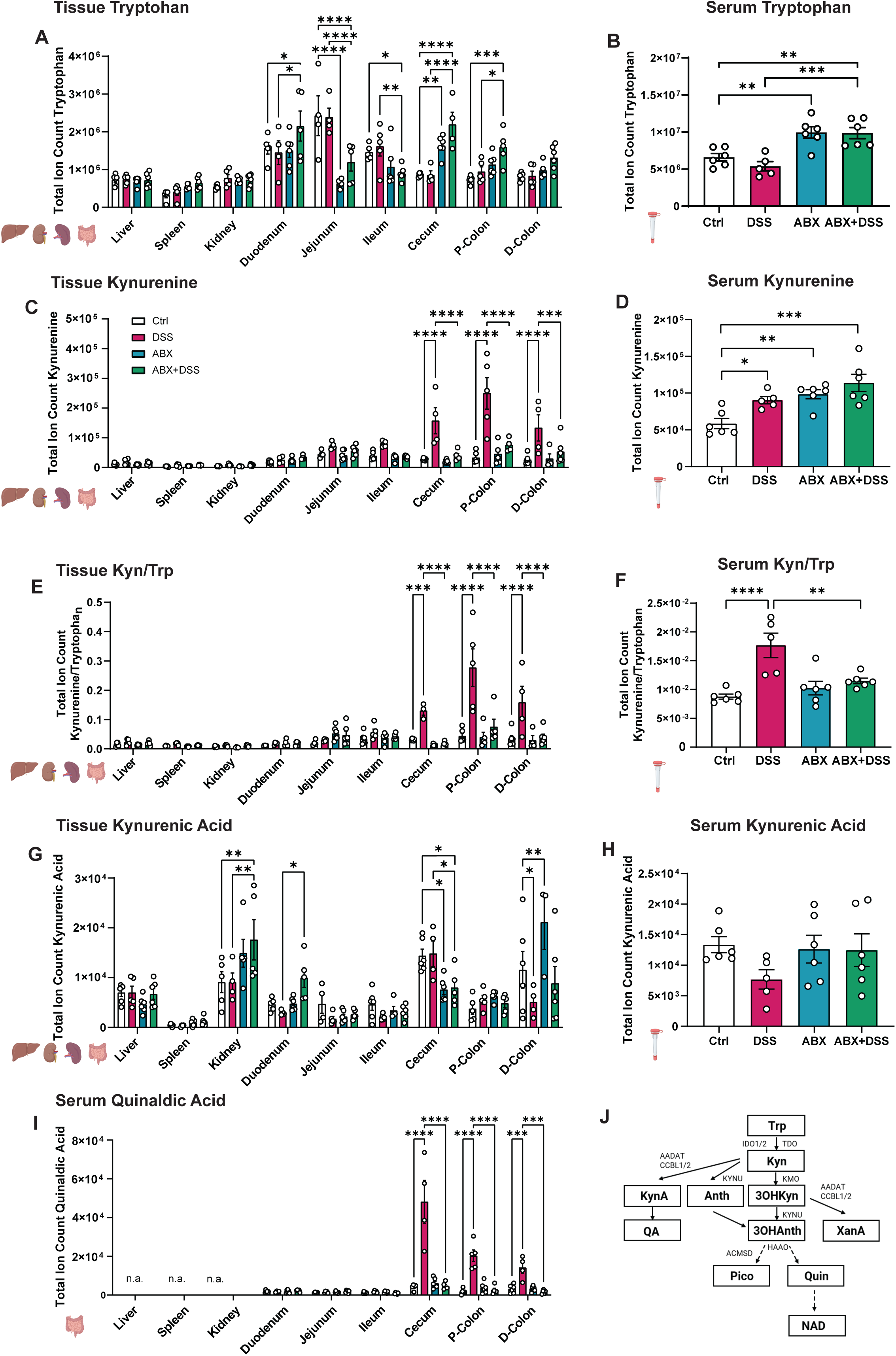
The gut microbiota influence turnover of Trp along the KP during DSS colitis. The indicated metabolites’ total ion counts were assessed in tissues (A,C,E,G,I) and serum (B,D,F,H). (J) KP overview. Data are presented as mean ± SEM. Statistical analysis was performed with two-way-ANOVA. ∗ p<0.05; ∗∗ p<0.01; ∗∗∗ p<0.001, ∗∗∗∗ p<0.0001. AADAT: kynurenine/alpha-aminoadipate aminotransferase, ACMSD: aminocarboxymuconate semialdehyde decarboxylase, CCBL: cysteine conjugate beta-lyase, HAAO: 3-hydroxyanthranilate 3,4-dioxygenase, IDO: indoleamine 2,3-dioxygenase, KMO: kynurenine-3-monooxygenase, KYNU: kynureninase, QPRT: quinolinate phosphoribosyltransferase, TDO2: tryptophan-2,3-Dioxygenase 2.

### The gut microbiota modulates the directionality of Trp catabolism along the KP during intestinal inflammation

Since ABX treatment increased Trp bioavailability in the host compartment, we next sought to determine how the intestinal microbiota drive host Trp degradation along the KP in the inflamed mucosa.

Pursuing the metabolic fate of Kyn degradation, we observed that increased Kyn levels in inflamed tissues (cecum, proximal/distal colon) of DSS-treated mice resulted in increased conversion into i) QA via kynurenic acid (KynA) (Fig. 3G-I), ii) XanA via 3-hydroxykynurenine (3OH-Kyn) (Fig. 4A,B) and iii) Quin via anthranilic acid (Anth), 3OH-Kyn and 3-hydroxyanthranilic acid (3OH-Anth) (Fig. 4,C-F). Levels of the intermediate catabolites KynA, 3OH-Kyn, Anth and 3OH-Anth remained largely unchanged (Fig. 3G,H, Fig. 4A,C,D). Corresponding to conversion of Trp via Kyn, 3OH-Kyn and 3OH-Anth into increased concentrations of Quin, gene expression of kynurenine 3-monooxygenase (*Kmo*) and 3-hydroxyanthranilate 3,4-dioxygenase (*Haao*) was upregulated, whereas quinolinate phosphoribosyltransferase (*Qprt*) was downregulated in comparison to control mice in the distal colon of DSS mice (Fig. S5B-D). The metabolic shift towards QA, XanA and Quin in inflamed intestinal tissue could be reverted in ABX-cotreated DSS colitis (Fig. 3I, Fig. 4B,E,F), coinciding with decreased expression of *Kmo* and *Haao* and restored expression of *Qprt* (Fig. S5B-D).

**Figure 4:**
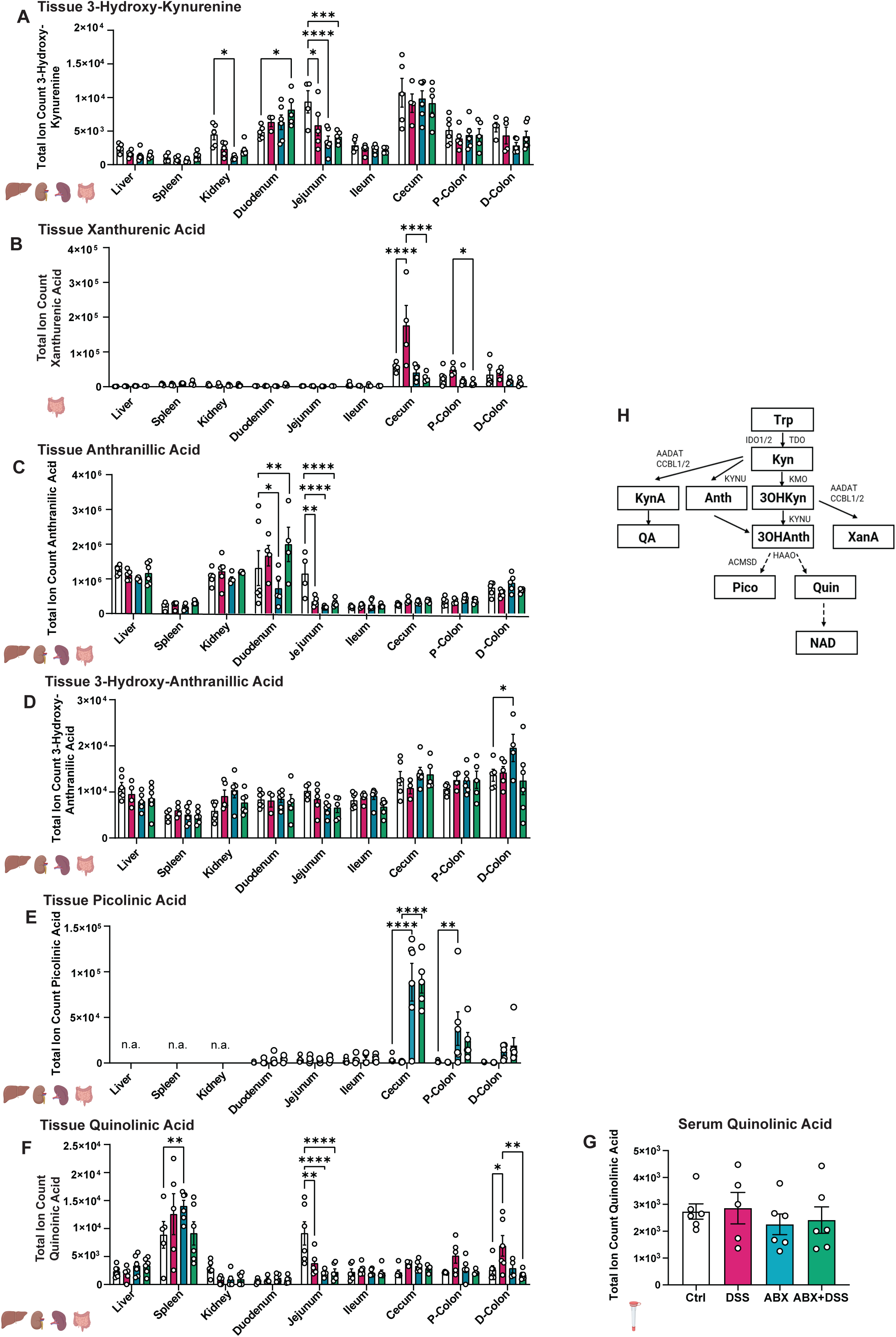
The gut microbiota modulate Trp metabolism through the KP during DSS colitis. The indicated metabolites’ total ion counts were measured in tissues (A-F) and serum (G). (H) KP overview. Data are presented as mean ± SEM. Statistical analysis was performed with two-way-ANOVA. ∗ p<0.05; ∗∗ p<0.01; ∗∗∗ p<0.001, ∗∗∗∗ p<0.0001. AADAT: kynurenine/alpha-aminoadipate aminotransferase, ACMSD: aminocarboxymuconate semialdehyde decarboxylase, CCBL: cysteine conjugate beta-lyase, HAAO: 3-hydroxyanthranilate 3,4-dioxygenase, IDO: indoleamine 2,3-dioxygenase, KMO: kynurenine-3-monooxygenase, KYNU: kynureninase, QPRT: quinolinate phosphoribosyltransferase, TDO2: tryptophan-2,3-Dioxygenase 2.

A remarkable shift was found in the conversion of 3OH-Anth into Quin or picolinic acid (Pico). Whereas the levels of 3OH-Anth were unaffected upon either DSS or ABX treatment in any assessed organ (Fig. 4D), we observed a striking dichotomy towards Pico in ABX-treated animals. Both Pico and Quin derive from the intermediate 2-amino-3-carboxymuconate-6-semialdehyde (ACMS), which can be converted into Pico by ACMS decarboxylase (ACMSD) or spontaneously form Quin [40]. Notably, Pico was not measurable in liver or kidney – organs with high ACMSD expression according to the Human Protein Atlas – whereas it accumulated specifically in the colonic mucosa, where *Acmsd* expression was undetectable across all experimental groups (data not shown). While we can rule out technical problems in Pico detection, we currently lack a clear biological explanation for this observation. Furthermore, DSS treatment amplified the previously described shift toward Quin accumulation in the inflamed mucosa (proximal and distal colon), which was reversed by ABX treatment (Fig. 4E,F). These findings largely coincided with metabolite concentrations within the luminal contents of the corresponding anatomical sites, despite Quin, which was decreased in luminal contents from DSS-treated mice (Figure S4C-G).

Overall, our findings suggest that in comparison to DSS-treated mice, only moderate abundances of Trp were degraded along an unobstructed KP in the colon of ABX-DSS mice. Furthermore, the directionality of Trp conversion in ABX-DSS mice shifted from accumulation of QA, XanA and Quin towards Pico, which implicates saturated flow into Quin and NAD^+^ *de novo* synthesis.

### Excess Trp upon microbial depletion rescues mucosal NAD(H) depletion during intestinal inflammation

Based on the hypothesis that increased Trp conversion along the KP reflects an attempt to sustain NAD⁺ *de novo* synthesis – and considering our previous finding of a metabolic bottleneck at QPRT within the inflamed colonic mucosa [27] – we next examined whether the metabolic changes observed in ABX-DSS mice resulted in restoration of NAD(H) metabolism. Indeed, NAD^+^ and NADH levels were recovered and partially even heightened in the colon of ABX-DSS mice (Fig. 5A,B). These findings coincided with rescued *Qprt* and nicotinamide nucleotide adenylyltransferase 3 (*Nmnat3*) expression in the distal colon (Fig. S5D,E), both enzymes responsible for converting Quin into NAD^+^. Notably, NAD⁺ and NADH levels were reduced in the luminal contents of ABX-DSS mice (Fig. 5C,D), suggesting that increased host Trp availability following microbial depletion and concurrent intestinal inflammation was redirected toward mucosal NAD(H) synthesis via restored metabolic flux through the kynurenine pathway. In contrast, an isolated increase in NAD(H) levels was detected in the luminal contents of the duodenum and jejunum of ABX-treated mice (Fig. 5C,D), suggesting that, under healthy conditions, microbes may normally consume NAD(H), whereas host cells – unlike in DSS-treated mice – are not in immediate need of it.

**Figure 5:**
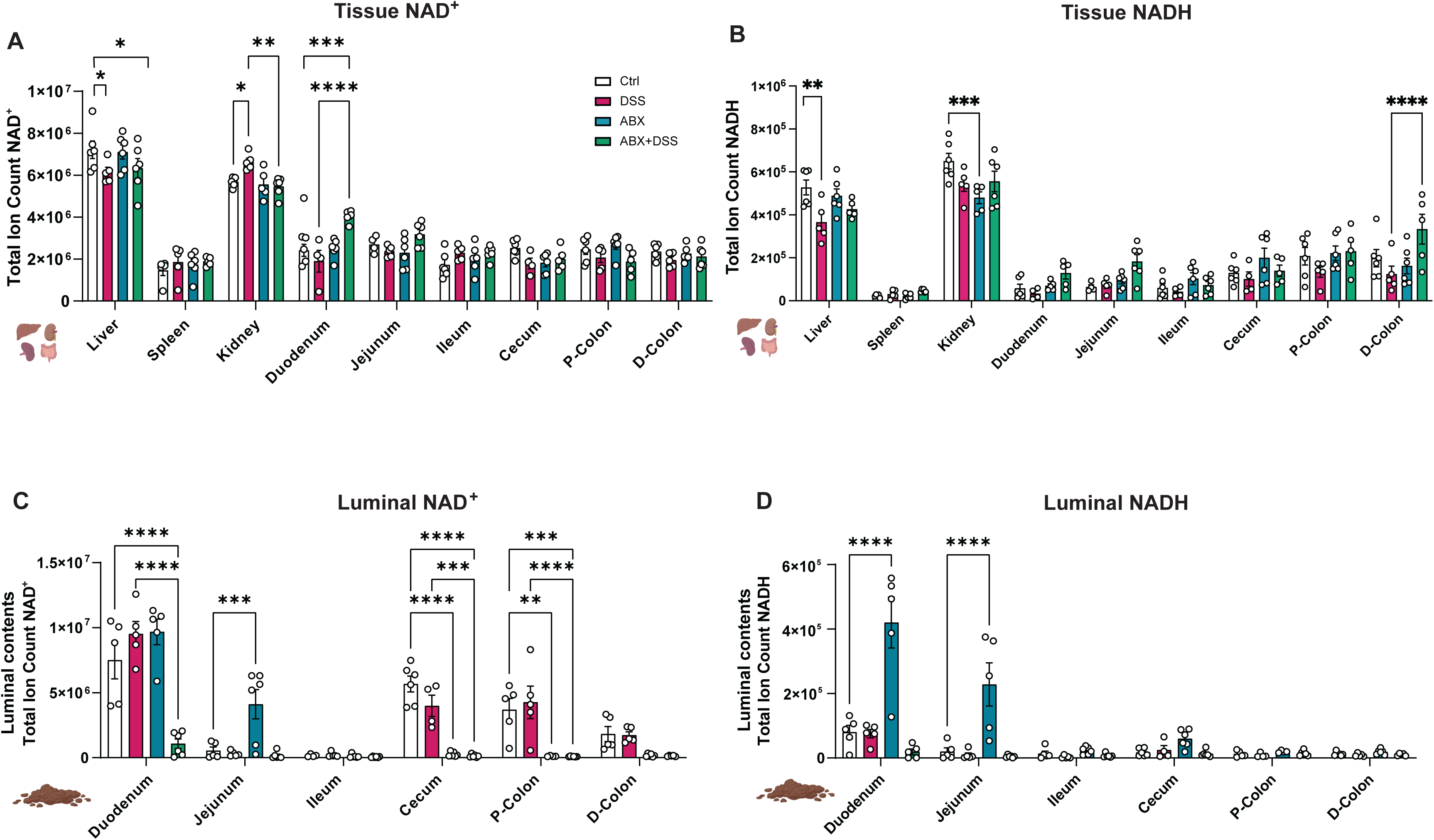
NAD(H) levels are influenced by DSS colitis and the gut microbiota. Total ion counts are shown for NAD^+^ and NADH in tissues (A,B) and luminal contents (C,D). Data are presented as mean ± SEM. Statistical analysis was performed with two-way-ANOVA. ∗ p<0.05; ∗∗ p<0.01; ∗∗∗ p<0.001, ∗∗∗∗ p<0.0001.

Furthermore, we unexpectedly observed that the metabolic alterations induced by microbial depletion were not confined to the inflamed colon but also occurred in the duodenum and jejunum: a significant increase of 3OH-Kyn, KynA and 3OH-Anth was observed in the duodenum of ABX-DSS mice, whereas abundances of these metabolites were reduced in the jejunum of ABX-DSS mice (Fig. 3G, Fig. 4A,D). These metabolic changes coincided with elevated abundances of Trp in the duodenum and reduced availability in the jejunum, respectively. Further intermediate catabolites of the KP remained unchanged within the duodenum and jejunum, however, augmented shuttling of Trp into the KP resulted in significantly elevated levels of NAD^+^ in the duodenum (Fig. 5A). Surprisingly, despite overall lowered abundances of KP precursor catabolites, we also observed heightened NADH abundances in the jejunum of ABX-DSS mice (Fig. 5B).

To assess whether restored mucosal NAD(H) levels in the colon of ABX-DSS mice might originate from other NAD(H) precursor pathways, such as the salvage pathway via NAM or NR, we measured NAM levels, which tended to be slightly reduced in GI tissues, luminal contents and serum of DSS- and DSS-ABX-treated mice, although no clear pattern was observed (Fig. 6A-C). However, gene expression of nicotinamide phosphoribosyltransferase (*Nampt*) was slightly elevated in the distal colon of ABX-treated mice (Fig. S5F), suggesting contribution of the salvage pathway to restored NAD(H) levels in the colon of ABX-DSS mice. Interestingly, microbial conversion of NAM into NA, which has previously been reported to fuel NAD^+^ metabolism in healthy mice [41,42], appeared to be disrupted in the colon of DSS mice (Fig. 6D). Accordingly, gene expression of nicotinate phosphoribosyltransferase (*Naprt*) was reduced in the distal colon of DSS and ABX-DSS mice (Fig. S5G). Overall, NAD(H) abundances were found to be higher in intestinal tissues from ABX-DSS mice than in tissues from control mice, offering a pathophysiological explanation for the amelioration of colitis severity in the respective mice.

**Figure 6:**
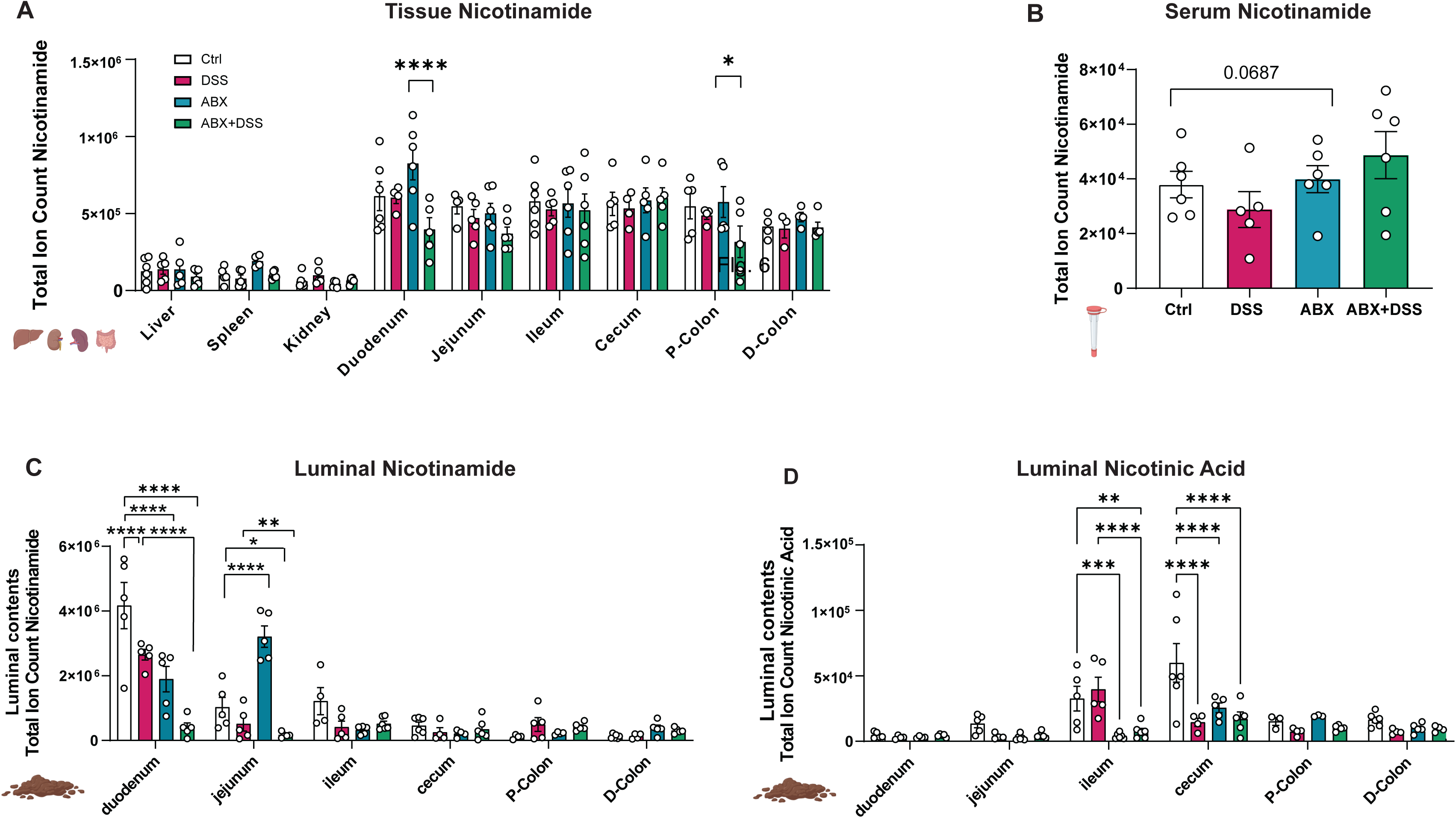
Antibiotics (pre)treatment modulates NAD^+^ precursors during DSS colitis. Total ion counts are demonstrated for NAM in tissues (A), serum (B) and luminal contents (C), NA is shown in luminal contents (D). Data are presented as mean ± SEM. Statistical analysis was performed with two-way-ANOVA. ∗ p<0.05; ∗∗ p<0.01; ∗∗∗ p<0.001, ∗∗∗∗ p<0.0001.

Having assessed how excess host Trp fuels the KP and NAD+ *de novo* synthesis, we lastly questioned whether it would also feed into the serotonin or indole pyruvate pathway. No corresponding changes in the levels of serotonin and indole in tissues and serum of ABX-DSS mice as compared to control or DSS-mice were observed (Fig. S6A,B,E). However, Trp promoted production of tryptamine in the duodenum, cecum, colon and serum (Fig. S6C,D), which was considered to be primarily host-driven, as microbially produced tryptamine was reduced in luminal contents of ABX-treated mice (Fig. S5H).

Of note, assessing gene expression of KP genes under physiologic conditions in intestinal tissues of GF mice and in mice held under specific pathogen free conditions, we found no significant changes. This suggests that upon immunologic equilibrium, host Trp metabolism is overall not markedly affected by the intestinal microbiota (Fig. S7). However, we have not conducted metabolic assessments of Trp-related metabolites in GF mice.

Taken together, our findings indicate that the elevated host Trp levels observed in inflammation-protected ABX-DSS mice result from a combination of reduced microbial consumption, enhanced absorption, and normalization of excessive Trp degradation through the KP. Dissecting the fate of excess Trp revealed that its conversion along the KP and subsequent NAD⁺ *de novo* synthesis were restored in the colon of ABX-DSS mice and markedly increased in the duodenum. Collectively, these results support a dual role for the gut microbiota during intestinal inflammation: (i) acting as a potent competitor for Trp degradation that limits host Trp bioavailability, and (ii) regulating systemic host homeostasis by shaping Trp and NAD^+^ metabolism.

## Discussion

The gut microbiota and immunometabolic processes at the host side are critically involved in IBD pathogenesis, but to which extend these processes are causally intertwined is poorly understood. Perturbed Trp and NAD^+^ metabolism have emerged as key metabolic phenomenons of chronic intestinal inflammation. We here conducted the first systemic, targeted analysis of Trp-related metabolites to understand how the intestinal microbiota shape Trp and NAD^+^ metabolism during intestinal inflammation induced by DSS as a preclinical model of colitis.

Depletion of the microbiota by a broad-spectrum ABX cocktail (metronidazole, vancomycin, neomycin and ampicillin) resulted in a marked reduction of bacterial abundances, consistent with the magnitude of depletion reported in previous studies using similar antibiotic regimens [43–45].

Reduction of the microbial load in line with several earlier reports significantly attenuated DSS-induced colitis; however, disease activity was not completely abolished, as indicated by a persistently elevated DAI and histopathological score at the end of the experiment (Fig. 1C,E). Of note, the phenotype of our ABX-DSS mice strongly deviated from the findings of Rakoff-Nahoum et al., whose study provided the basis for the antibiotic cocktail used here. While they reported aggravated DSS-induced colitis upon microbial depletion, our ABX-DSS-treated mice exhibited attenuated colitis. Several factors may account for this discrepancy, including differences in mouse strains, housing conditions, or DSS type and regimen (2% for 7 days in their study vs. 2.5% for 5 days; similar molecular weight) [46]. These differences potentially contributed to the fact that, unlike in our experiments, DSS-treated wildtype mice in the study of Rakoff-Nahoum et al. did not exhibit a similarly pronounced weight loss, which biases phenotype-based comparisons to ABX-DSS-treated animals across experiments [15].

While amelioration of colitis phenotypes following depletion of the gut microbiota – as observed here – has been reported in several studies, opposing effects have been described as well. These divergent outcomes imply context-dependent interactions between host and gut microbiota, wherein microbiota may act as either protectors or drivers of intestinal inflammation. Protective effects are likely mediated by reduced microbiota-driven immune activation – for instance, through impaired expansion of colitogenic bacteria or enrichment of beneficial genera that produce anti-inflammatory metabolites such as short-chain fatty acids (SCFA) or modulate mucus structure [47–49]. Detrimental effects of microbial depletion have on the other hand been attributed to a disruption of intestinal barrier function and epithelial integrity, impaired establishment of mucosal immune cell tolerance, or loss of interactions between toll-like receptors (TLRs) and commensal bacteria, which physiologically promote epithelial homeostasis and barrier integrity [11,15,50].

We here observed that during DSS-induced colitis, gut bacteria either produced or positively influenced the production of several metabolites with known anti-inflammatory properties in feces and tissues: Next to SCFA, these metabolites included Kyn, XanA, QA, tryptamine, IPA or ILA, of which the first four have been described to originate both from host and the gut microbiota, whereas production of IPA and ILA is microbe-specific (Fig. 2B-D, G,H, Fig. 3C,D,I, Fig. 4B) [51–54]. Similarly, synthesis of these beneficial metabolites was hampered upon ABX co-treatment, underlining the crucial role of the microbiota in regulating their formation (Fig. 2B-D, G,H). Anti-inflammatory effects of these compounds are among other mechanisms exerted via the aryl hydrocarbon receptor (AhR), or via direct restoration of mitochondrial function in cell types involved in the immune responses contributing to IBD pathogenesis [55–58]. While altered fecal metabolite patterns could result from accelerated intestinal transit and increased luminal water content, overt diarrhoea could be excluded, as stool consistency was only mildly affected in all groups. These findings indicate that the gut microbiota contributes to counteracting murine intestinal inflammation through the generation of anti-inflammatory catabolites.

Depletion of gut microbiota increased Trp abundances not only in feces and in the luminal compartment, but also in serum and tissues (Fig. 2A,N, Fig. 3A,B) [59,60]. Several physiological processes might explain this observation, including i) reduced microbial Trp degradation and ii) diminished mucosal inflammatory burden, resulting in a lowered demand of Trp degradation along the KP for *de novo* NAD^+^ synthesis. While we observed microbial production of protective Trp-derived metabolites such as SCFA, tryptamine, IAA and IPA during DSS colitis, IBD-related dysbiosis disrupts the production of those anti-inflammatory Trp catabolites [61–63]. This may indicate unique metabolic responses to either acute or chronic intestinal inflammation, possibly influenced by differing microbial compositions and their metabolic outputs. From a therapeutic perspective, it may be beneficial to selectively introduce bacterial strains or their metabolites that support Trp-related protective mechanisms in IBD.

Following the fate of elevated Trp in mice cotreated we DSS and ABX, we found a striking difference when comparing DSS to DSS-ABX animals. Whereas in DSS-challenged mice, we observed reduced mucosal NAD(H), ABX co-treatment not only normalised the passage of Trp through the KP but also restored mucosal NAD(H) levels (Fig. 5A,B). Of note, we observed these effects where predominantly host-driven, supported by the near-total depletion of NAD(H) derivatives in feces and luminal contents of ABX-DSS mice (Fig. 2I-M, Fig. 5C,D). This finding was unexpected, since the gut microbiota has been reported to relevantly contribute to host NAD(H) metabolism; however, increased host Trp utilization following microbial depletion allowed it to compensate and even enhance NAD(H) levels [41,64]. Furthermore, reduced fecal NAM abundances have been associated with dysbiosis in UC, suggesting the microbial capacity to support NAD⁺ synthesis is impaired during colitis [65].

Despite reduced levels of anti-inflammatory catabolites (IPA, SCFA, QA, XanA) in ABX-DSS-mice, mucosal restocking of NAD(H) constitutes a novel, compelling metabolic explanation for the reduced inflammatory activity in those mice. Emphasizing the importance of intact NAD^+^ synthesis for the resolution of inflammation, we and others have shown that supplying other NAD^+^ precursors than Trp (NR, NAM) rescues murine intestinal inflammation [27,29]. Translating these preclinical observations, an oral controlled-ileocolonic-release formulation of NAM is currently tested for its efficacy in mild-moderate UC (CICR-NAM; NCT06488625). Consistently, we observed faster recovery of COVID-19 patients when supplementing NAM [66]. We here embark on these findings by providing evidence that mucosal *Qprt* expression and thus, NAD^+^ *de novo* synthesis, is critically regulated by gut microbiota during intestinal inflammation, opening a so far undescribed therapeutic window for targeted restoration of depleted mucosal NAD^+^ levels.

An evident limitation of our study is the difficulty of disentangling directionality: do alterations in Trp and NAD^+^ metabolism mediate the amelioration of inflammation upon microbial depletion, or are these metabolic shifts a secondary consequence of anti-inflammatory effects conferred by microbial depletion? Nevertheless, our data indicate that changes in Trp metabolism – e.g., increased host Trp bioavailability – already emerge under ABX treatment alone and become further aggravated when DSS-induced inflammation is superimposed. Notably, excess Trp upon microbial depletion in DSS-treated mice was actively shuttled into the KP and efficiently fuelled NAD⁺ *de novo* synthesis despite an insufficiently resolved inflammatory phenotype (Fig. 1), generating a metabolic profile clearly distinct from control and DSS-mice. A direct way to further evaluate this would be to compare DSS versus ABX-DSS mice under conditions of Trp starvation or pharmacological or genetic blockade of the KP at Ido1 or Qprt, which would allow to determine whether NAD⁺ *de novo* synthesis causally contributes to ameliorated inflammation. Another limitation of our study is the exclusive use of the DSS-induced colitis model. While this epithelial injury-driven model does not capture the full complexity of human IBD, it is an acknowledged, robust and reproducible system to study microbiota-host metabolic interactions during acute colonic inflammation, making it well suited for the mechanistic insights presented here [67].

In summary, we show that diminishing the microbiota during DSS colitis impairs bacterial Trp catabolism, thereby enhancing host Trp access. This metabolic shift restores mucosal NAD⁺ *de novo* synthesis, which we propose as a novel pathomechanism of alleviated intestinal inflammation upon microbial depletion.

## Declarations

### Ethics approval and consent to participate

All animal experiments were either performed at the Pennsylvania State University and approved by the Institutional Animal Care and Use Committee (IACUC - PROTO202202188).) or at the Central Animal Facility (ZTH) of the University Hospital Schleswig Holstein (UKSH, Kiel, Germany) approved by the local animal safety review board of the federal ministry of Schleswig Holstein (IX 554 - 62160/2024 (48-6/24) and internal §4 project number 1410).

## Consent for publication

This study did not involve human data.

## Availability of data and materials

Raw metabolomics data were generated at Pennsylvania State University, PA, US, and deposited at the Zenodo repository (https://zenodo.org/communities/abxdss-metabolomics/), linking to all datasets. Each dataset is also individually accessible through the following accession links https://doi.org/10.5281/zenodo.17500534, https://doi.org/10.5281/zenodo.17500751, https://doi.org/10.5281/zenodo.17500765, https://doi.org/10.5281/zenodo.17500761. Sequencing analyses were not conducted in this study. No special codes were utilized.

## Acknowledgments

We gratefully acknowledge the excellent work and support of our colleagues Vivian Sauer and Jaedon Sadler. We would further like to acknowledge the Huck Institutes’ Metabolomics Core Facility (RRID:SCR_023864) for use of the OE 240 LC-MS and Sergei Koshkin for helpful discussions on sample preparation and analysis.

## Author Contributions

K.A., M.R.M., P.R., L.W., A.I.A, D.M.M.H. designed the study.

A.I.A., L.W., D.M.M.H., M.Y., T.M., F.S., J.B., E.S., S.W. performed experiments and analyzed the data.

K.A., P.R., L.W., A.I.A, D.M.M.H. planned the project and supervised the experiments. K.A., L.W. wrote the initial manuscript.

K.A., S.W., M.R.M., P.R., S.Schr. edited the manuscript

## Abbreviations

Ahr: Aryl hydrocarbon receptor
3OH-Anth: 3-hydroxy-anthranilic acid
3OH-Kyn: 3-hydroxy-kynurenine
Anth: Anthranilic acid
CD: Crohn’s disease
DSS: dextran sodium sulfate
HIAA: 5-hydroxy indoleacetic acid
IAA: indole-3-acetic acid
IBD: Inflammatory bowel disease
IDO1/2: Indoleamine 2,3-dioxygenase
1/2 ILA: Indolelactic acid
IPA: Indole-3-propionic acid
KP: Kynurenine pathway
KMO: Kynurenine-3-monooxygenase
Kyn: Kynurenine
KynA: Kynurenic acid
KYNU: Kynureninase
NA: Nicotinic acid
NAD+: Nicotinamide adenine dinucleotide
NAM: Nicotinamide
NAPRT: Nicotinic acid phosphoribosyltransferase
NMNAT3: Nicotinamide nucleotide adenylyltransferase 3
NR: Nicotinamide riboside
Pico: Picolinic acid
QPRT: Quinolinate phosphoribosyltransferase
Quin: Quinolinic acid
QA: quinaldic acid
TDO: Tryptophan-2,3-dioxygenase
Trp: Tryptophan
UC: Ulcerative Colitis
XanA: Xanthurenic acid

---

**Figure S1:**
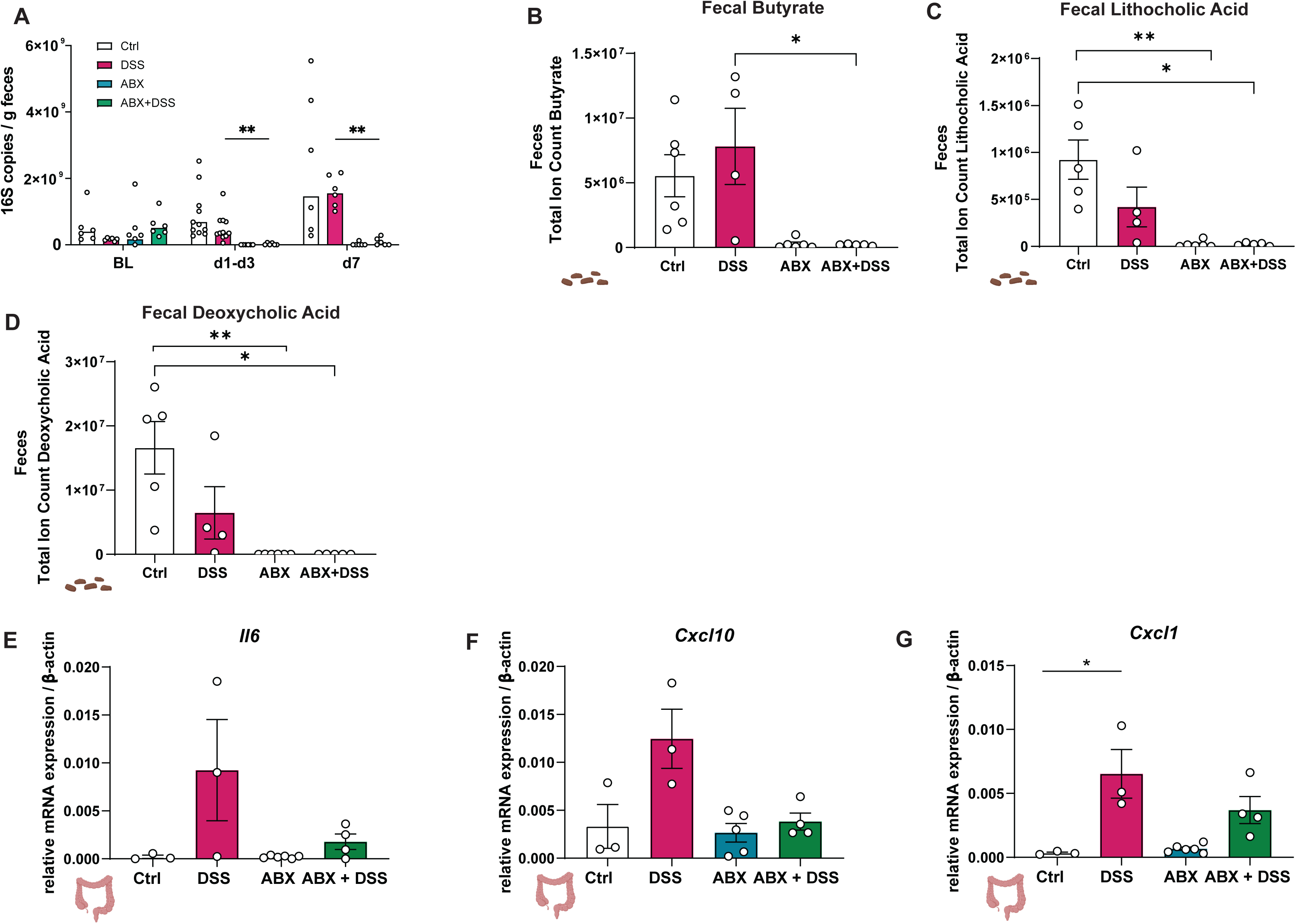
Antibiotics (pre)treatment effectively depletes the gut microbiota. (A) 16S rRNA gene copy number was quantified by qPCR to assess bacterial abundance. Mixed-effects analysis with Šídák’s post-hoc test was used for statistical analysis. (B-D) Total ion counts were measured for the indicated metabolites. (E-G) RT-qPCR was conducted from colonic tissue (distal colon) for the indicated cytokines. β-actin served as a housekeeping gene. Data are presented as mean ± SEM. Statistical analysis was performed with two-way-ANOVA. ∗ p<0.05; ∗∗ p<0.01.

**Figure S2:**
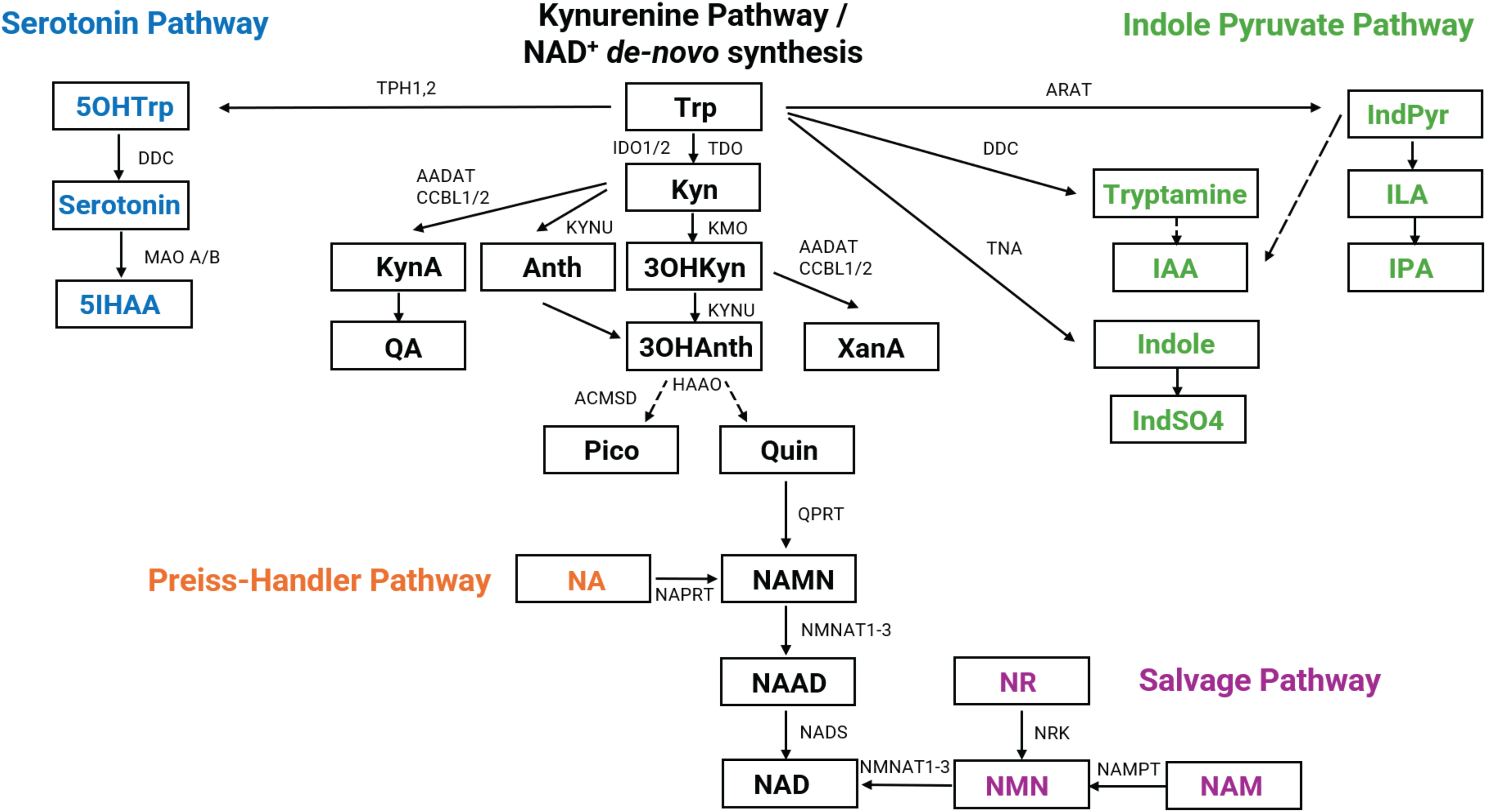
Overview over Trp-degrading pathways. Trp is degraded along the kynurenine pathway (KP) (black), the serotonin pathway (blue) or the indole pyruvate pathway (green). NAD^+^ can be synthesized by i) *de novo* synthesis from Trp, ii) via the Preiss-Handler Pathway from NA or iii) via the salvage pathway from NAM or NR. AADAT: kynurenine/alpha-aminoadipate aminotransferase, ACMSD: aminocarboxymuconate semialdehyde decarboxylase, Anth: anthranilic acid, ARAT: aromatic amino acid aminotransferase, 3OH-Anth: 3-hydroxy-anthranilic acid, CCBL: cysteine conjugate beta-lyase, HAAO: 3-hydroxyanthranilate 3,4-dioxygenase, DDC: dopa decarboxylase, IAA: indoleacetic acid, IDO: indoleamine 2,3-dioxygenase, ILA: indolelactic acid, IndSO4: indoxyl sulfate, IndPyr: indole-3-pyruvate, IPA: indolepropionic acid, 5HIAA: 5-hydroxy indoleacetic acid, KMO: kynurenine-3-monooxygenase, Kyn: kynurenine, KynA: kynurenic acid, KYNU: kynureninase, 3OH-Kyn: 3-hydroxy-kynurenine, NA: nicotinic acid, NAAD: nicotinic acid adenine dinucleotide, NAD: nicotinamide adenine dinucleotide, NADS: nicotinamide adenine dinucleotide synthetase 1, NAM: nicotinamide, NAMN: nicotinic acid mononucleotide, NAMPT: nicotinamide phosphoribosyltransferase, NAPRT: nicotinate phosphoribosyltransferase, NMN: nicotinamide mononucleotide, NMNAT: nicotinamide nucleotide adenylyltransferase, NR: nicotinamide riboside, NRK: nicotinamide riboside kinase, Pico: picolinic acid, QA: quinaldic acid, Quin: quinolinic acid, QPRT: quinolinate phosphoribosyltransferase, TDO2: tryptophan-2,3-dioxygenase 2, TNA: tryptophanase, TPH: tryptophan hydroxylase, Trp: tryptophan, XanA: xanthurenic acid, 5OH-Trp: 5-hydroxy-tryptophan.

**Figure S3:**
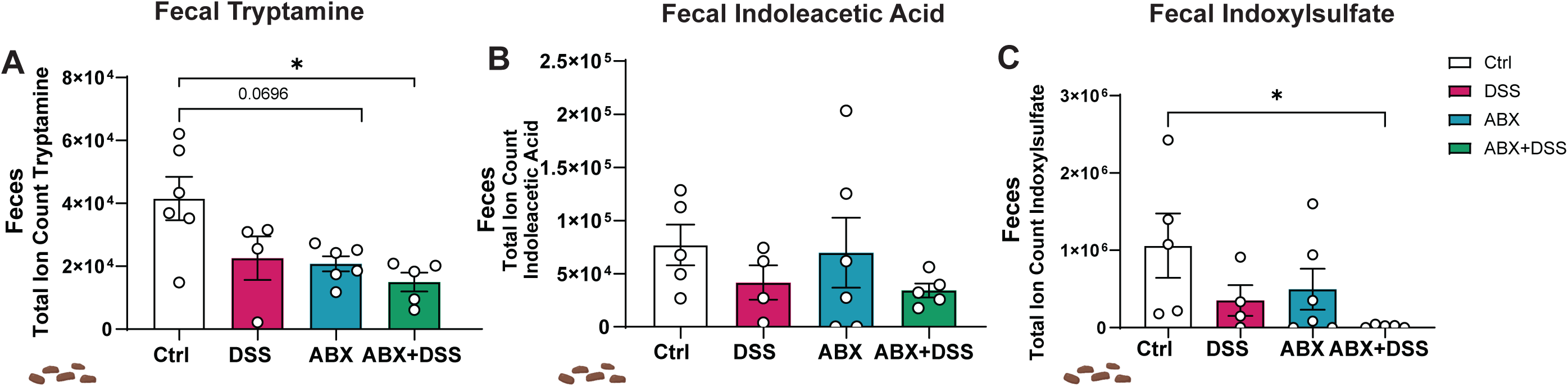
Fecal metabolites are altered by DSS colitis and antibiotics (pre)treatment. (A-C) Total ion counts of the indicated metabolites are shown in feces. Data are presented as mean ± SEM. Statistical analysis was performed with two-way-ANOVA. ∗ p<0.05.

**Figure S4:**
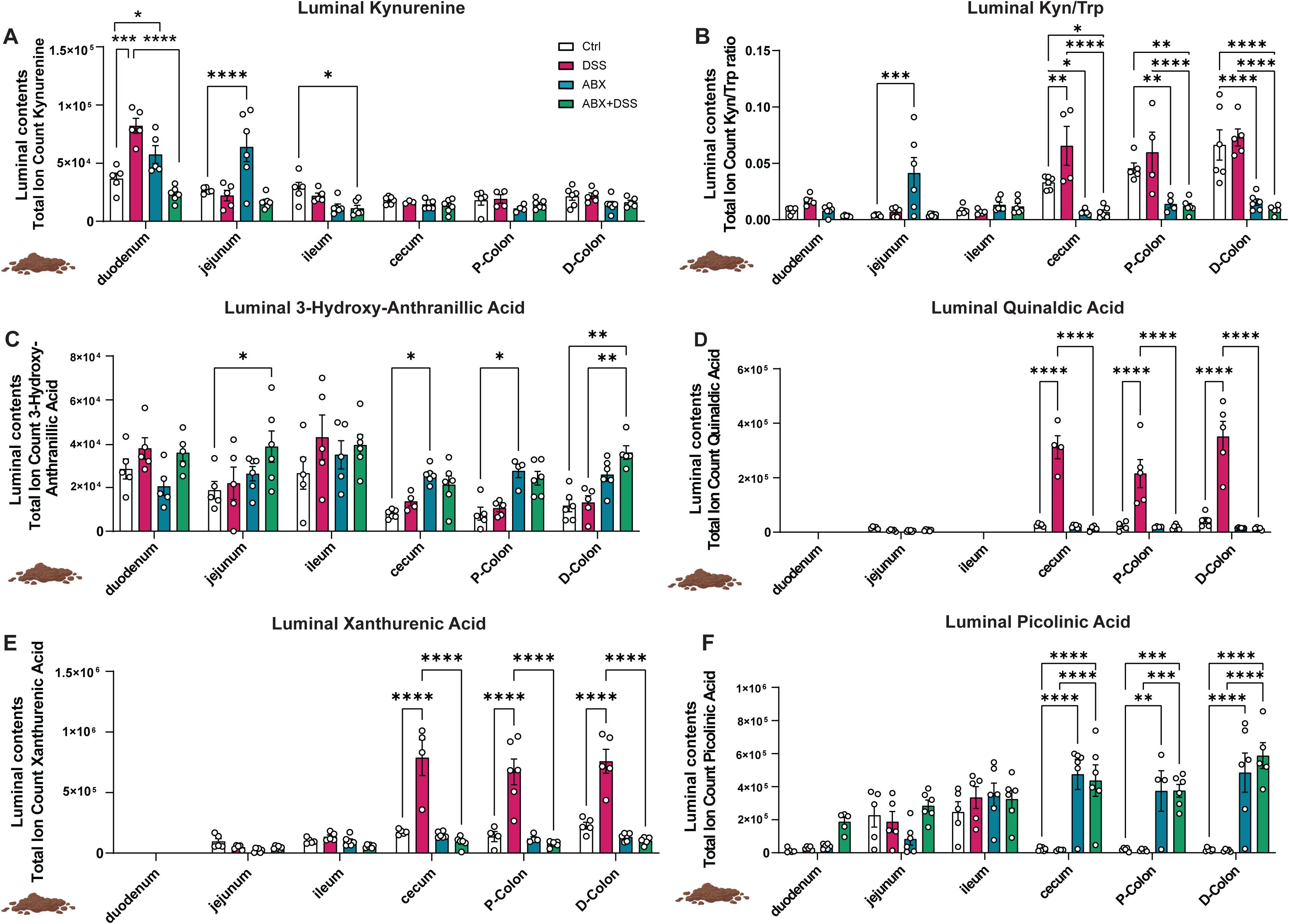
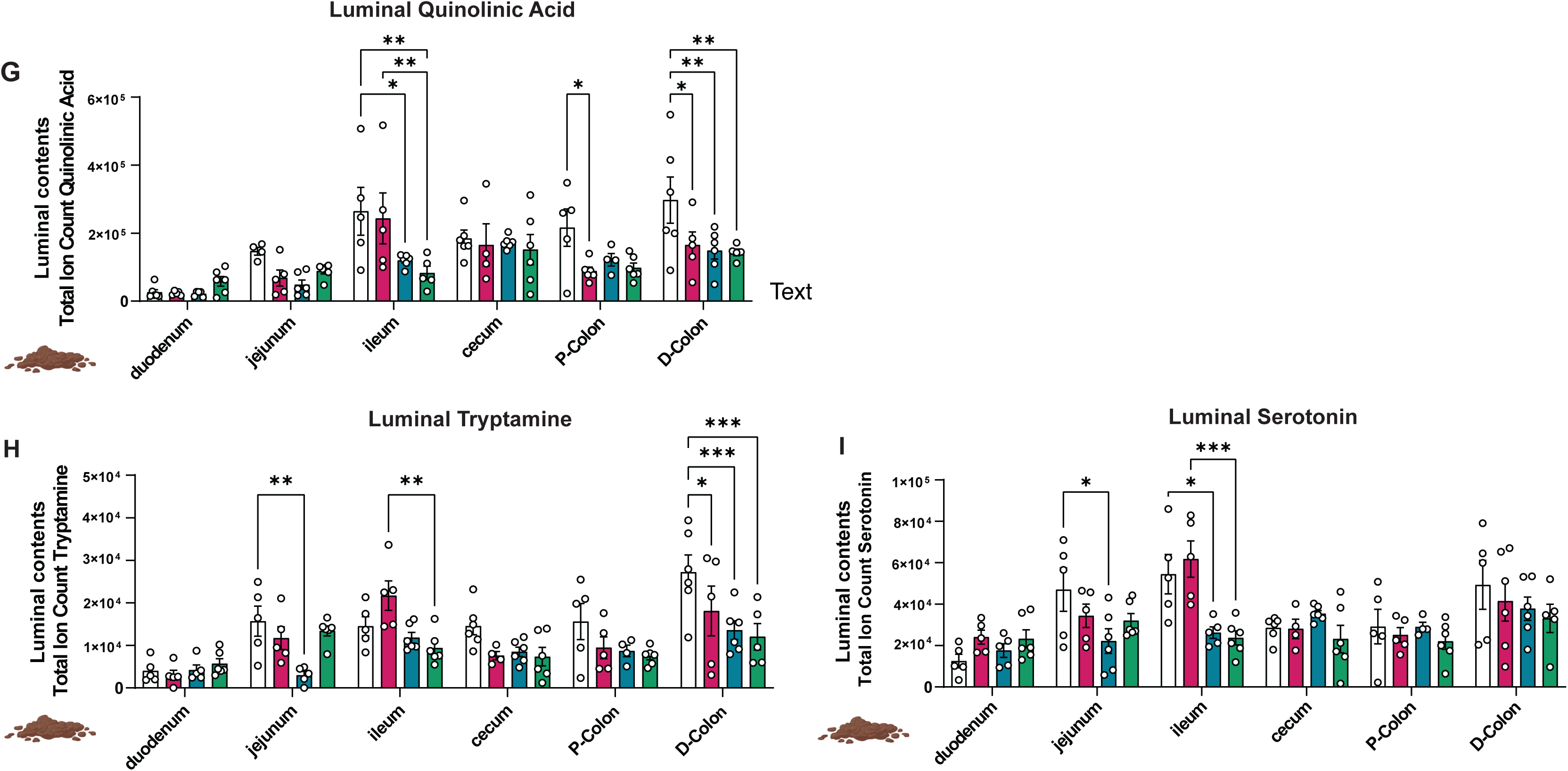
The intestinal microbiota modulate host Trp degradation along the KP during gut inflammation. (A-I) Total ion counts of the indicated metabolites were measured in luminal contents. Data are presented as mean ± SEM. Statistical analysis was performed with two-way-ANOVA. ∗ p<0.05; ∗∗ p<0.01; ∗∗∗ p<0.001, ∗∗∗∗ p<0.0001.

**Figure S5:**
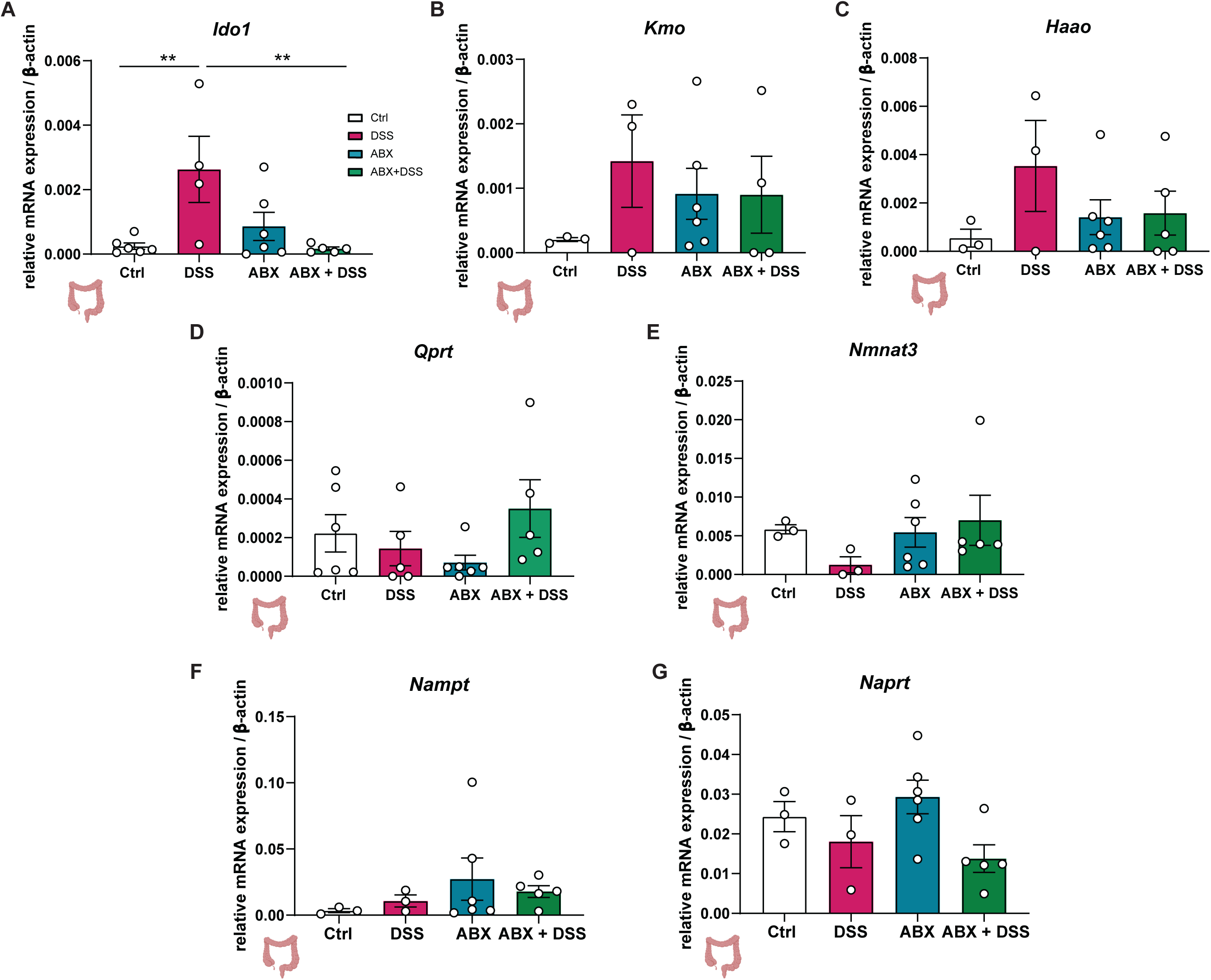
Host colonic gene expression is altered by microbial depletion and DSS colitis. (A) RT-qPCR was conducted from colonic tissue (distal colon) for the indicated KP genes. β-actin served as a housekeeping gene. Data are presented as mean ± SEM. Statistical analysis was performed with two-way-ANOVA. ∗∗ p<0.01. Ido1: indoleamine 2,3-dioxygenase 1, Haao: 3-hydroxyanthranilate 3,4-dioxygenase, Kmo: kynurenine 3-monooxygenase, Nampt: nicotinamide phosphoribosyltransferase, Naprt: nicotinate phosphoribosyltransferase, Nmnat3: nicotinamide nucleotide adenylyltransferase 3, Qprt: quinolinate phosphoribosyltransferase.

**Figure S6:**
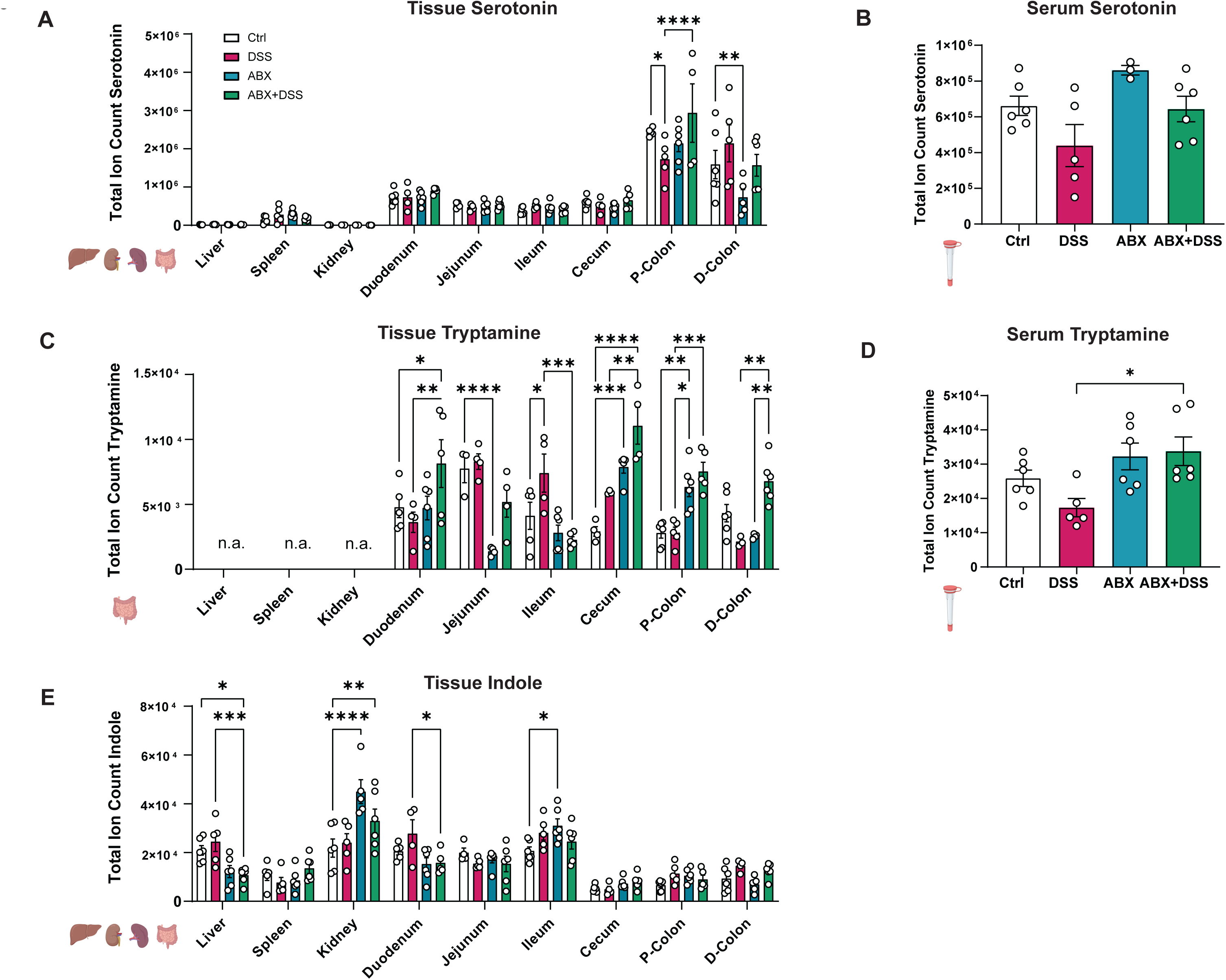
Tryptamine levels increase upon DSS colitis and antibiotics (pre)treatment. Total ion counts of the indicated metabolites were assessed in tissues (A,C,E) and in serum (B,D) Data are presented as mean ± SEM. Statistical analysis was performed with two-way-ANOVA. ∗ p<0.05; ∗∗ p<0.01; ∗∗∗ p<0.001, ∗∗∗∗ p<0.0001.

**Figure S7:**
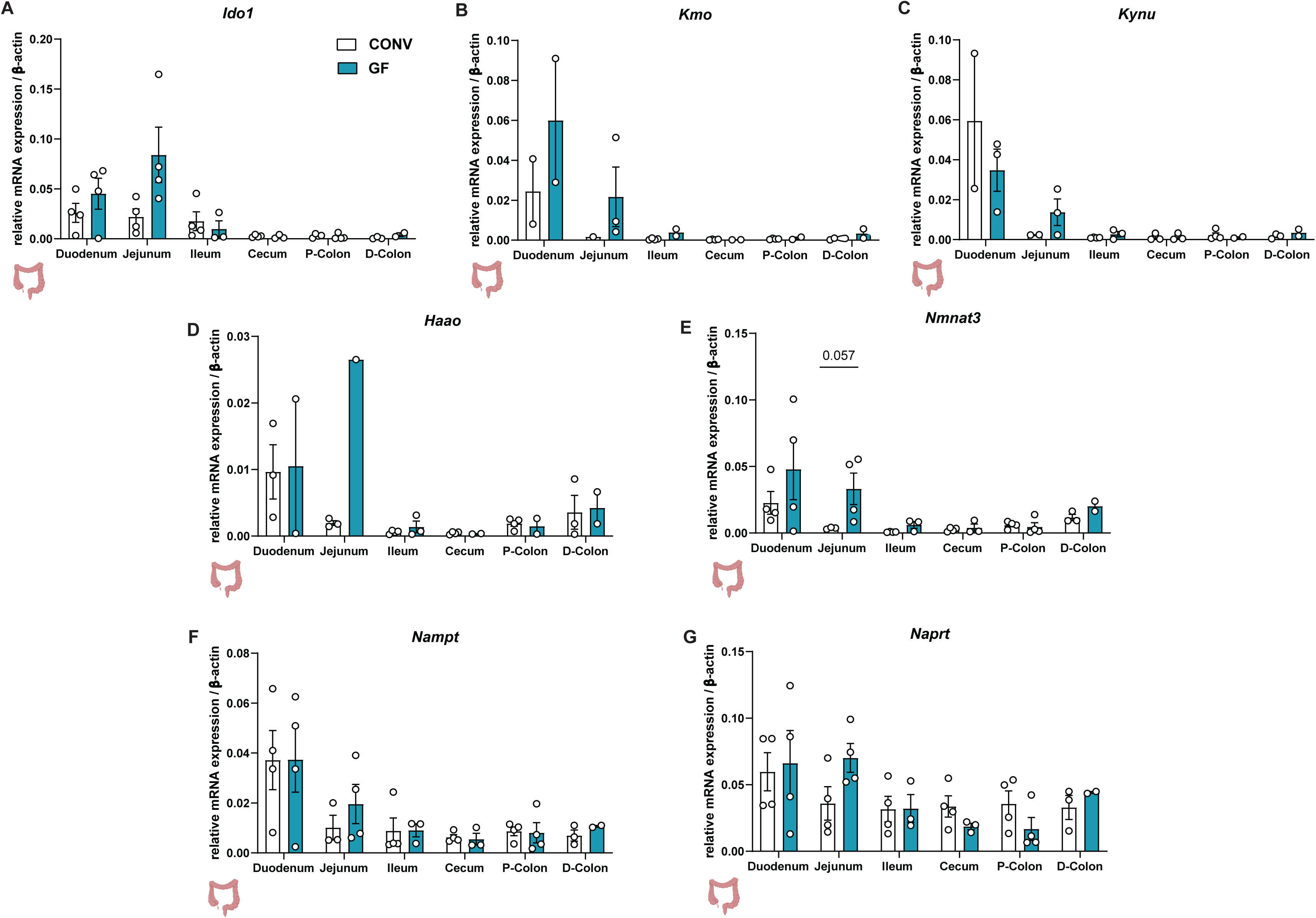
Microbial depletion alone does not affect host colonic gene expression without intestinal inflammation. (A) RT-qPCR was performed with tissue derived from the distal colon for the indicated KP genes in control (CONV) and germ-free (GF) mice. β-actin served as a housekeeping gene. Data are presented as mean ± SEM. Statistical analysis was performed with Mann-Whitney U test. Ido1: indoleamine 2,3-dioxygenase 1, Haao: 3-hydroxyanthranilate 3,4-dioxygenase, Kmo: kynurenine 3-monooxygenase, Kynu: kynureninase, Nampt: nicotinamide phosphoribosyltransferase, Naprt: nicotinate phosphoribosyltransferase, Nmnat3: nicotinamide nucleotide adenylyltransferase 3.

